# Identification of 4,5,6,7-Tetrabromo-1*H*-benzotriazole (TBB) as a Small Molecule MESH1 Inhibitor that Suppresses Ferroptosis

**DOI:** 10.64898/2026.02.19.706832

**Authors:** Alexander A. Mestre, Yunju Oh, Jianli Wu, Denise Dunn, Yasaman Setayeshpour, Ssu-Yu Chen, Chao-Chieh Lin, C. Skyler Cochrane, Pyeonghwa Jeong, Gibeom Nam, Chloe Markey, Daniel Reker, Scott R. Floyd, Jiyong Hong, Pei Zhou, Jen-Tsan Chi

**Affiliations:** Department of Biochemistry, Duke University School of Medicine, Durham, NC, 27710, USA; Department of Molecular Genetics & Microbiology, Duke University School of Medicine, Durham, NC, 27710, USA; Department of Chemistry, Duke University, Durham, NC, 27708; Department of Radiation Oncology, Duke University School of Medicine, Durham, NC, 27710, USA; Department of Cell & Molecular Biology, Duke University School of Medicine, Durham, NC, 27710, USA; Department of Biomedical Engineering, Duke University, Durham, NC, 27710, USA; Department of Pharmacology and Cancer Biology, Duke University School of Medicine, Durham NC, 27710, USA; Center for Advanced Genomic Technologies, Duke University, Durham NC, 27710, USA

**Keywords:** MESH1, HDDC3, Ferroptosis, NADPH, TBB, 4, 5, 6, 7-tetrabromo-1H-benzotriazole

## Abstract

Ferroptosis is a regulated form of cell death driven by iron-dependent lipid peroxidation and contributes to diverse pathologies including ischemia-reperfusion injury and neurodegenerative disorders. Current ferroptosis inhibitors largely function as nonspecific radical-trapping antioxidants, limiting their clinical utility. We previously identified MESH1 as a key regulator of ferroptosis through its NADPH phosphatase activity. Here, we identify 4,5,6,7-tetrabromo-1*H*-benzotriazole (TBB) as a small molecule inhibitor of MESH1 with an IC_50_ value of 4.7 ± 0.3 µM. X-ray crystallography revealed the molecular determinants of TBB recognition which are corroborated through structure-activity relationships of TBB analogs. TBB protected multiple cell lines against ferroptosis *in vitro*, and this effect was mitigated by *MESH1* knockdown, consistent with on-target activity. Furthermore, TBB reduced neuronal death in an *ex vivo* brain slice model of Alzheimer’s disease. Collectively, these findings establish TBB as a *bona fide* small-molecule MESH1 inhibitor that suppresses ferroptosis and establishes MESH1 as a promising therapeutic target.

**Graphical Abstract:** Depicting mechanism of TBB suppressing ferroptosis through the inhibition of MESH1. Figure Created with Biorender.com

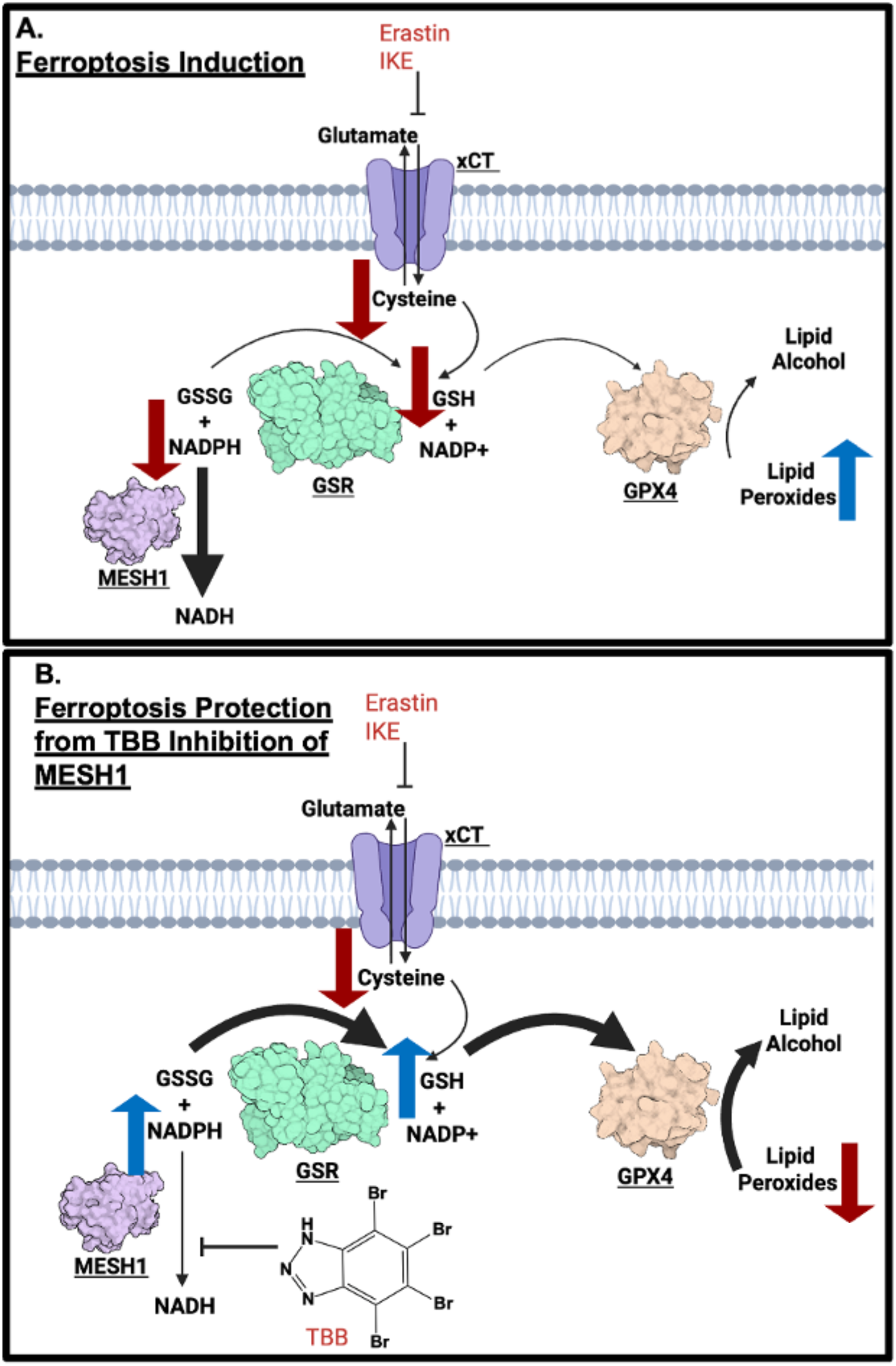

## Introduction

Ferroptosis is a form of regulated cell death, which is characterized by iron accumulation and lipid peroxidation (1). Ferroptosis was discovered during the study of how the small molecule erastin triggered cell death (2). Subsequent studies demonstrated that erastin inhibits the cysteine/glutamate antiporter (system X_C-_), leading to a non-apoptotic form of cell death termed “ferroptosis” (3). There is growing evidence that ferroptosis plays an important role in diverse human diseases, including cancer, acute kidney disease, metabolic dysfunction-associated steatotic liver disease (MASLD), neurodegeneration, stroke, and ischemic-reperfusion injury (4–8). Over the last decade, there has been an explosion of research that has started to unravel the cellular mechanisms and molecular determinants regulating ferroptosis. Collectively, these studies have converged on a critical role of metabolic controls, with pathways regulating thiols, lipids, and iron, lying at the core of ferroptosis (9).

All organisms face continuous metabolic stress and require coping mechanisms to ensure survival and maintain homeostasis. In bacteria, the primary metabolic stress response is the stringent response, which is regulated by the alarmone (p)ppGpp. This system triggers transcriptional reprogramming that enhances survival while limiting growth (10). The intracellular levels of (p)ppGpp are regulated by the RelA synthetase and the SpoT hydrolase. Metazoan genomes contain a SpoT homolog called MESH1 (<u>Me</u>tazoan <u>S</u>poT <u>H</u>omologue <u>1</u>) that is capable of hydrolyzing (p)ppGpp and regulating stress response genes in *Drosophila* (11). However, metazoan genomes lack any gene encoding a (p)ppGpp synthetase, and the physiological relevance of the low (p)ppGpp levels in animal cells remains unclear (12, 13). Consequently, the endogenous substrates and functions of MESH1 in metazoans remained unclear until our discovery of MESH1 as an NADPH phosphatase that converts NADPH to NADH (14). In parallel, we recently identified an additional role for MESH1 acting as a phosphatase toward PAPS, revealing a distinct role for MESH1 in sulfur metabolism (15). NADPH serves as a pivotal metabolite within the ferroptosis-associated metabolic pathways, and its intracellular concentration functions as a biomarker of ferroptosis sensitivity (16). Functionally, NADPH is used to regenerate the critical antioxidant glutathione (GSH). We discovered that the NADPH phosphatase activity of MESH1 regulates ferroptosis and that genetic knockdown of *MESH1* protects cells against ferroptosis by preserving NADPH levels (14). Since our initial discovery, the NADPH phosphatase activity of MESH1 homologues in *C. elegans* (17) and bacteria (18) has been independently verified, indicating that this function is evolutionarily conserved across kingdoms. Furthermore, the role of MESH1 in mediating ferroptosis has also been reported in the contexts of neonatal cardiomyopathies (19) and hepatocellular carcinoma (20). Together, these findings establish MESH1 as a conserved regulator of ferroptosis and have motivated our investigation of small-molecule MESH1 inhibitors as potential ferroptosis-targeted therapeutics.

There is growing interest in developing therapeutics targeting ferroptosis. While most efforts have focused on ferroptosis inducers in the context of cancer research, the mounting evidence that ferroptosis contributes to neurodegeneration and ischemic injury has spurred the parallel interest in developing ferroptosis inhibitors. Ferroptosis inhibitors have been categorized into several categories, including radical trapping antioxidants (RTAs), lipoxygenase (LOX) inhibitors, and iron chelators (21). RTAs consist of endogenous radical-trapping antioxidants such as Vitamin E (alpha-tocopherol), coenzyme Q10, and Vitamin K, and synthetic RTAs such as ferrostatin, which has become the reference compound for ferroptosis inhibition (21). Although both endogenous and synthetic RTAs potently suppress lipid peroxidation *in vitro,* they encounter fundamental mechanistic and translational limitations: because each RTA molecule can neutralize only a finite number of lipid radicals, achieving physiological relevance requires exceedingly high concentrations (22, 23). This intrinsic constraint likely underlies the repeated failure of endogenous and synthetic RTAs to demonstrate clinical benefit in numerous large-scale trials (24). Another category of ferroptosis inhibitors includes LOX (lipoxygenase) inhibitors such as FDA-approved zileuton, which can inhibit LOX enzymes isoforms, responsible for the lipid peroxidation of polyunsaturated fatty acids (PUFAs), however, can also act as an RTAs (25, 26). Finally, iron chelators such as deferoxamine (DFO) can inhibit ferroptosis by chelating cytosolic iron and/or the iron in the active site of LOX enzymes (3).

Since MESH1 is essential for ferroptotic cell death by acting as an NADPH phosphatase and since its removal confers ferroptosis protection, we reasoned that small-molecule inhibitors of MESH1 could provide a pharmacologic route to ferroptosis suppression. By preserving intracellular levels of NADPH, MESH1 inhibitors should maintain the reducing power needed to support endogenous enzymatic antioxidant mechanisms (e.g., glutathione- and thioredoxin-dependent systems) that counter lipid peroxidation and regenerate antioxidant capacity. This mechanism is distinct from RTAs and does not have the same stoichiometric limitations as RTAs. Here, we present our discovery and biochemical, structural, and functional characterization of TBB as a MESH1-targeting ferroptosis inhibitor identified through screening of a small molecule kinase library, thereby establishing MESH1 as a viable therapeutic target for ferroptosis-driven diseases.

## Results

### Screening Kinase Inhibitor Library

Although MESH1 is a phosphatase, its engagement with nucleotide substrates (ppGpp, NADPH) suggests that its binding pocket resembles the ATP-binding sites of kinases. Accordingly, we screened the Comprehensive Kinase Screening Library (Cayman), comprising ∼850 compounds, to identify inhibitors of MESH1 **(Figure 1A)**. Phosphate release from NADPH hydrolysis was quantified by the malachite green assay as described previously (14). Each compound was tested at a single concentration of 10 μM; reactions were incubated for 10 min and then quenched. For each compound the percentage of MESH1 activity was normalized against the DMSO control, and compounds were ranked by inhibitory potency **(Figure 1B)**. Four compounds reduced MESH1 activity by more than 50%. Among the top hits were 4,5,6,7-tetrabromo-1H-benzotriazole (TBB) and three catechol-containing natural products: piceatannol, hispidin, and butein. As catechol-containing compounds are frequently flagged as Pan Assay Interference Compound (PAINS) that can inhibit targets through multiple nonspecific mechanisms (27), we prioritized TBB for further characterization.

**Figure 1.**
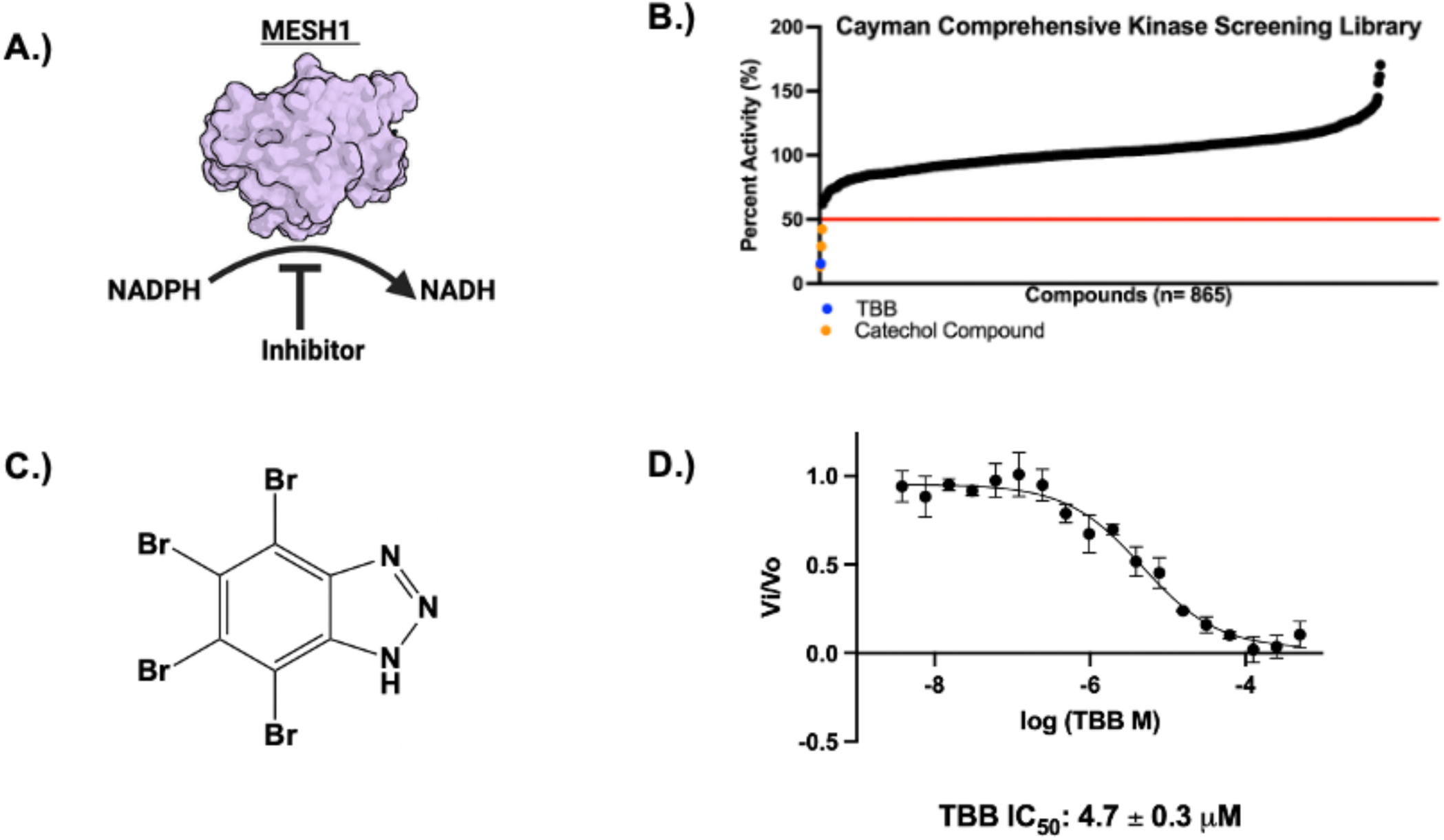
**A.** Graphical representation of inhibitor screen strategy monitoring MESH1 NADPH phosphatase activity. **B**. High throughput screen of kinase library. Each compound ranked by potency with compounds that reduced percent activity below 50% (red line) were classified as hits. TBB shown as blue dot, catechol containing compounds were shown with orange dots, all other compounds were shown as black dots (n=1). **C.** Chemical structure of TBB **D.** IC50 curve of TBB fitted with four-parameter fitting (n= 3 independent experiments, mean ± S.D.)

TBB consists of a brominated benzotriazole architecture that mimics the base of a nucleoside **(Figure 1C)**. Its IC_50_ value was calculated from an 18-point dose response curve **(Figure 1D)**. TBB had an IC_50_ value of 4.7 ± 0.3 µM and a K_i_^app^ of 2.3 ± 0.2 µM under the mode of competitive inhibition **(Figure 1D)**. TBB is a selective inhibitor (IC_50_ of 0.3 µM and a K_i_ of 0.64 µM) of the alpha subunit of Casein Kinase-2 (CK2), a constitutively active serine/threonine kinase involved in apoptosis suppression (28, 29). Accordingly, the K_i_^app^ value of 2.3 ± 0.2 µM places MESH1 among the high-affinity cellular targets of TBB and suggests that previous studies attributing phenotypes solely to CK2 inhibition might inadvertently include a significant contribution from MESH1 inhibition.

### Structural analysis of the MESH1-TBB Complex

To elucidate the molecular details of the MESH1-TBB interaction, we determined the crystal structure of the MESH1-TBB complex by soaking TBB into apo MESH1 crystals. The MESH1-TBB crystals diffracted to a resolution of 2.33 Å **(Figure 2A; Supplemental Table 1).** In the MESH1-TBB structure, MESH1 had little conformational change and adopted a architecture of ten α-helices similar to that in previously solved MESH1 structures in the apo (PDB: 3NR1, RMSD: 0.199 Å) and NADPH-bound (PDB: 5VXA, RMSD: 0.306 Å) states **(Figure 2A)** (11, 14).

**Figure 2.**
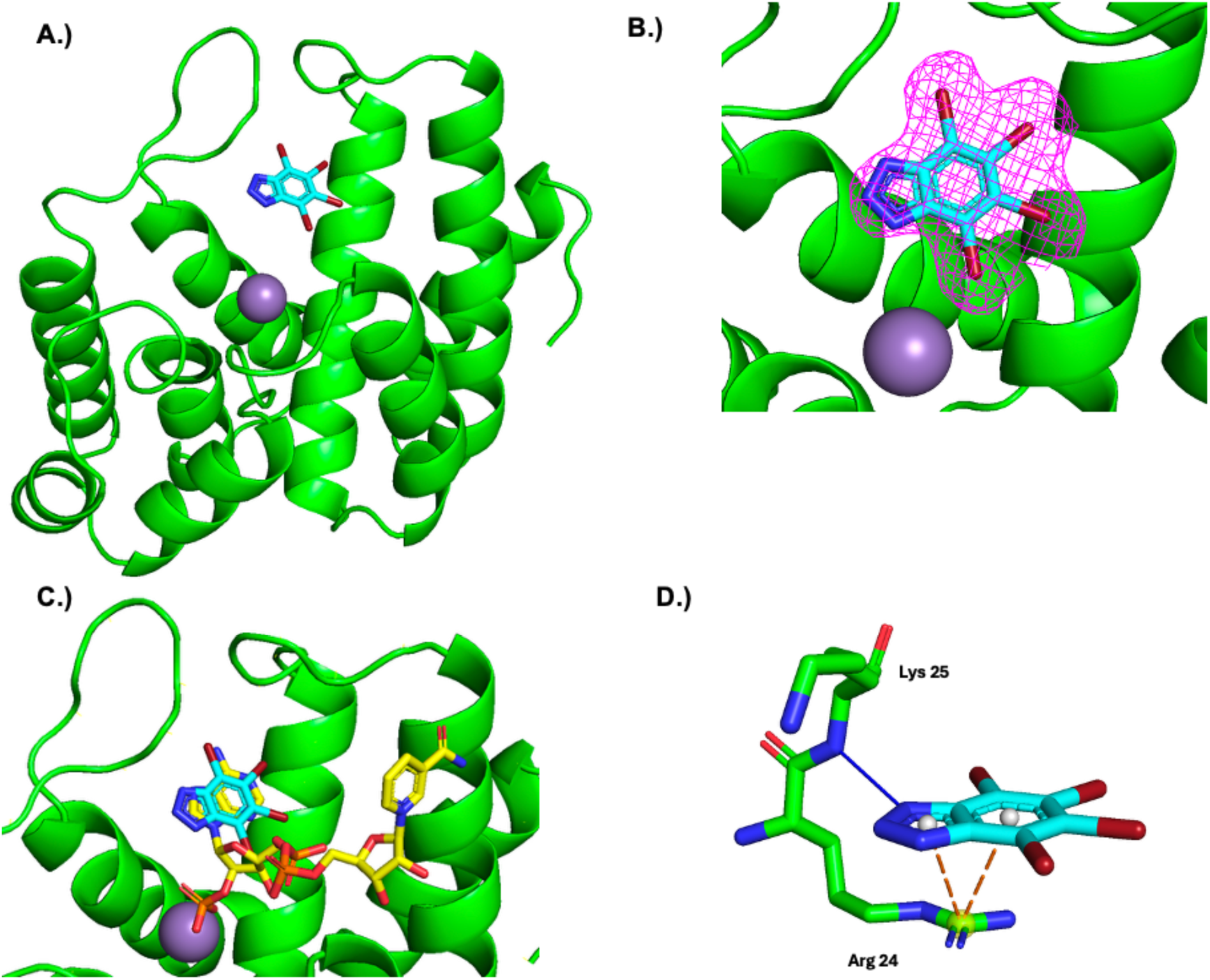
**A.** TBB bound to the active site of MESH1. MESH1 is shown in green ribbon diagram. TBB shown as blue stick model. Mn^2+^ shown as grey sphere. **B.** 2mFo-DFc map of the density corresponding to TBB contoured to 𝛔1. **C.** Comparing the binding sites of TBB (PDB: 9ZZ9, shown as blue stick model) and NADPH (PDB: 5VXA shown as yellow sticks) **D.** Interactions between MESH1 and TBB. Hydrogen bond shown as solid blue line. π-cation interaction shown as orange dashed line.

The active site of MESH1 comprises a substrate-binding pocket that accommodates nucleotide substrates, positioned above an HD motif that coordinates a catalytic Mn²⁺ ion essential for phosphatase activity **(Figure 2A)**. The 2mFo-DFc omit map reveals well-defined electron density for TBB within the nucleotide-binding site **(Figure 2B)**, where TBB adopts a binding pose similar to the adenine base of NADPH **(Figure 2C)**. TBB makes several interactions with MESH1 **(Figure 2D)**. The 3-nitrogen forms a hydrogen bond with the amide group of Lys-25; the same interaction occurs between the Lys-25 amide and the 7-nitrogen of NADPH (14). Additionally, Arg-24 is positioned to form a π-cation interaction with the extended aromatic system of TBB. The 4-bromine inserts into a small cavity that is normally occupied by the exocyclic N6-amine on the adenosine moiety of NADPH. The 5- and 6-bromines of TBB face toward the hydrophobic region of the nucleotide binding pocket, where they contribute to hydrophobic interactions. The 7-bromine is the most solvent exposed facing away from the nucleotide binding site towards the rest of the open active site. TBB buries approximately 324 Å^2^ of its 370 Å^2^ solvent-accessible surface area upon binding, consistent with a hydrophobic molecule engaging a partially enclosed binding pocket. Overall, the crystal structure reveals how TBB binds to MESH1 and provides a structural framework for interpreting its inhibitory activity that could inform future efforts for lead optimization.

The TBB-MESH1 interaction shares similarities with the TBB-CK2α interaction (30). Like the MESH1 binding site, the CK2α ATP binding site consists of a narrow hydrophobic cavity. However, unlike the single primary conformation of TBB binding to MESH1, TBB adopts many different binding conformations when binding to CK2α fluctuating between a tug-of-war between hydrophobic effect/ halogen bonding with the backbone carbonyls of Glu-114 and Val-116, and electrostatic interaction/ hydrogen bond with Lys-68 (30). The additional halogen bonds and electrostatic interactions may help explain why TBB has ∼3.5 fold increased selectivity for CK2 (K_i_: 0.64 µM) in comparison to MESH1 (K_i_^app^: 2.3 µM) (28).

### Structure-Activity Relationships of TBB Analogs

To determine the structure-activity relationship (SAR) between the TBB chemical structure and MESH1 inhibition, we analyzed a series of TBB analogs **(Figure 3)**. These analogs were studied using orthogonal approaches, including enzyme inhibition (percent activity) and thermal shift (ΔTm) at the inhibitor concentration of 100 µM. These orthogonal approaches produced highly correlated results **(Supplemental Figure 1A)**. Additional IC_50_ values were measured for compounds that displayed significant inhibition and stabilization.

**Figure 3.**
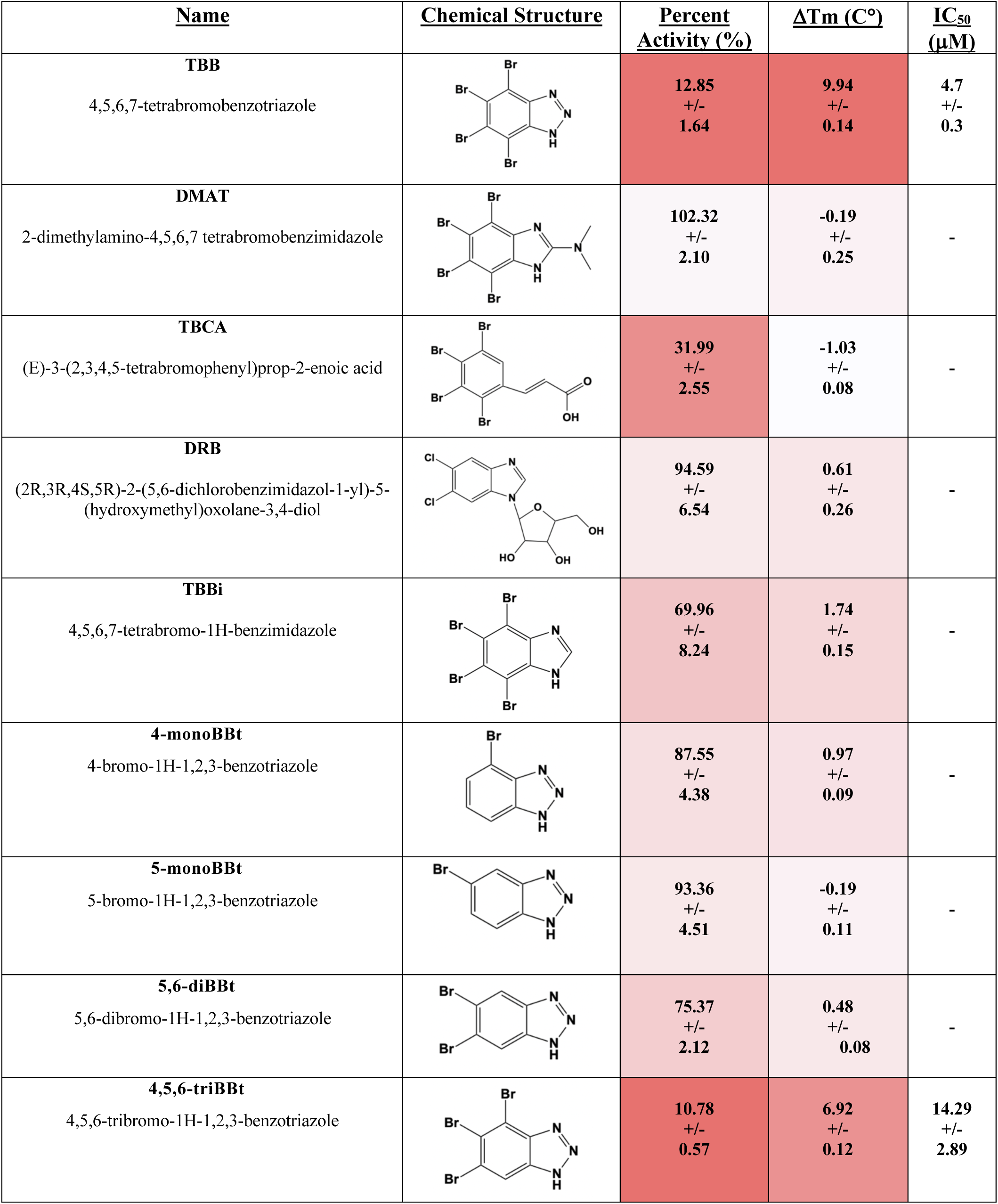
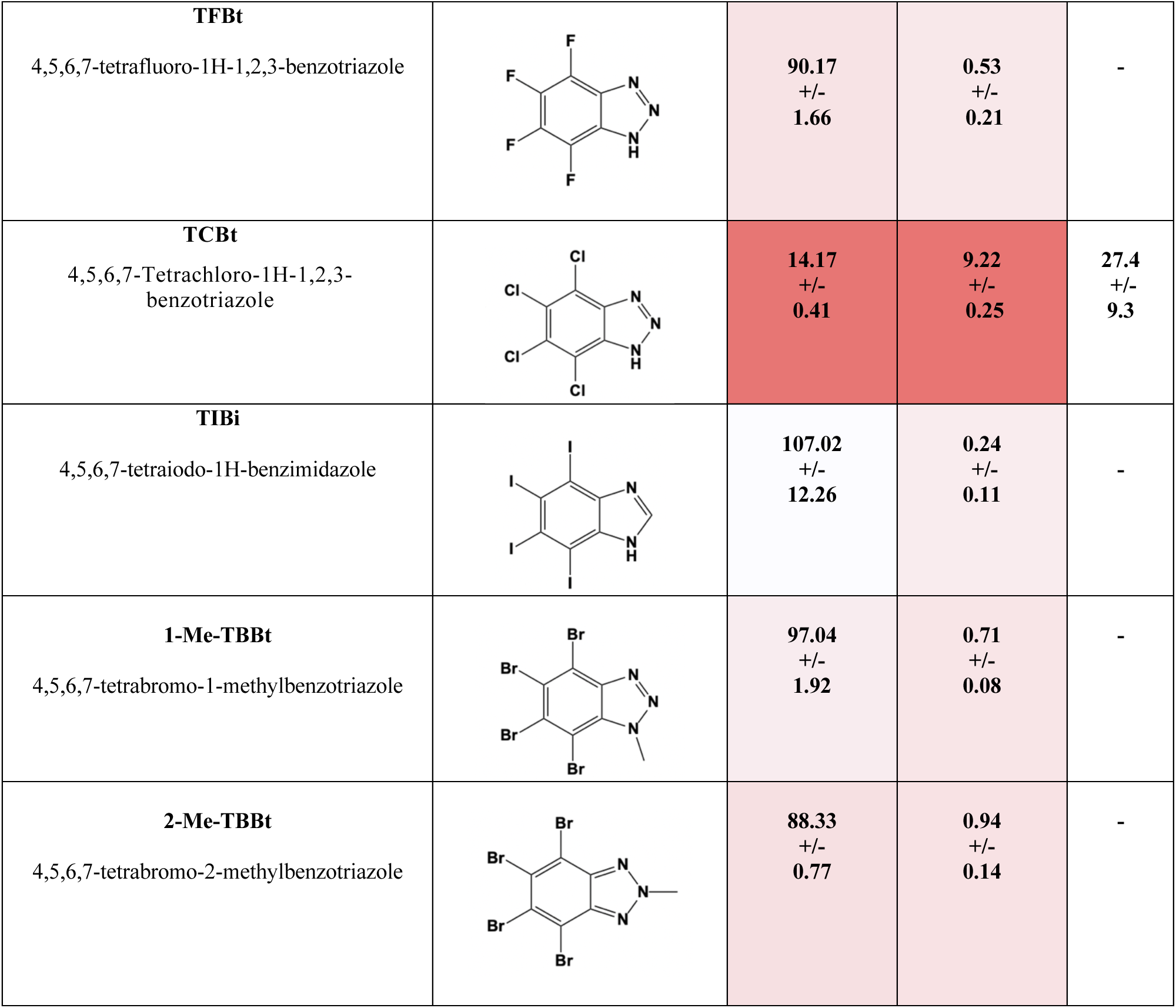
SAR of TBB analogs tested. Heat map red color = stronger inhibition and larger thermal shift and white = weaker inhibition and smaller thermal shift.

There has been considerable effort to develop TBB analogs with increased specificity towards CK2 (31, 32). Some of the most historically important TBB analogs include 2-dimethylamino-4,5,6,7-tetrabromo-1*H*-benzimidazole (DMAT), (*E*)-3-(2,3,4,5-tetrabromophenyl)prop-2-enoic acid (TBCA), and (2*R*,3*R*,4*S*,5*R*)-2-(5,6-dichlorobenzimidazol-1-yl)-5-(hydroxymethyl)oxolane-3,4-diol (DRB). DMAT replaces the triazole ring with a tertiary amine-substituted imidazole ring, TBCA eliminates the triazole ring entirely and replaces it with an n-alkenyl carboxylate group, whereas DRB substitutes the triazole ring with a ribose-attached imidazole ring and additionally replaces the tetrabromo substituents with dichloro atoms. None of these CK2-inhibiting TBB analogs inhibited or stabilized MESH1, highlighting major differences between the MESH1 and CK2 active sites.

To investigate the significance of the 1,2,3-triazole group, we examined the CK2 inhibitor 4,5,6,7-tetrabromo-1*H*-benzimidazole (TBBi), which has a single atom change from 2-nitrogen to carbon, replacing the 1,2,3-triazole ring with an imidazole. This single-atom change resulted in a drastic decrease in inhibition (∼6-fold decrease in percent activity), underscoring the significance of the triazole functional group for TBB to inhibit MESH1.

Next, we investigated how the number of bromine substitutions influenced the activity of TBB. We tested single bromine variants at the 4 and 5 positions of the benzene (4-monoBBt and 5-monoBBt). Additionally, an analog doubly brominated at the 5 and 6 positions of the benzene (5,6-diBBt) was tested. Finally, we synthesized a triple-bromine analog bearing substitutions at the 4,5, and 6 positions (4,5,6-triBBt). These bromine variants revealed the importance of multiple bromine substitutions, showing a linear relationship between the number of bromines and inhibition/stabilization **(Supplemental Figure 1B)**. The single and double bromine analogs failed to inhibit MESH1, while the triple brominated analog, 4,5,6-triBBt, displayed an IC_50_ of 14.29 ± 2.89µM, representing a ∼3-fold weaker inhibition in comparison to TBB **(Supplemental Figure 1C)**.

We also examined the impact of different halogenated analogs on MESH1 inhibition. We tested full 4,5,6,7 substitutions of the bromines of TBB with fluorides and chlorides. The 4,5,6,7-fluorine substitution (TFBt) failed to inhibit or stabilize MESH1. The 4,5,6,7 chlorine-substituted molecule (TCBt) had similar inhibition and ΔTm to TBB at 100 µM inhibitor, and the IC_50_ value of TCBt was calculated to be 27.4 +/- 9.3 µM, which is ∼6-fold weaker than TBB **(Supplemental Figure 1D)**. Intriguingly, TCBt is a known inhibitor for the West Nile Virus Helicase with an IC_50_ value of 27 µM, making MESH1 one of the better molecular targets for TCBt (33). It has been reported that 4,5,6,7-tetraiodo-benzotriazole cannot be synthesized (34). Instead, we compared 4,5,6,7-tetraiodo-1H-benzimidazole (TIBi) with TBBi and found that the fully iodinated phenyl substitution resulted in weaker inhibition than TBBi. These results indicate that complete iodination at these bromine positions reduced TBB activity. These results indicate that for uniform halogen substitutions at the 4,5,6,7 positions, potency increased along with the size of the halogen atom F < Cl < Br. However, the SAR data also indicated that Br was optimal, as iodine substitution decreased inhibition.

The triazole moiety of TBB has an experimentally measured pKa of 4.78, corresponding to the deprotonation of the neutral triazole ring to form an anionic species at physiological pH (35).

Deprotonation of the triazole group has previously been shown to be important for CK2 inhibition (35). To assess whether this deprotonation state is also required for MESH1 inhibition, we synthesized TBB analogs that methylated the N1 or N2 positions of the triazole group (1-Me-TBBt and 2-Me-TBBt), thereby preventing deprotonation and locking the molecule into a neutral state. Unlike TBB, neither of these methylated compounds inhibited or stabilized MESH1, indicating that a negative charge is essential for MESH1 inhibition. This requirement also explains the weaker activity of TBBi, which contains an imidazole ring that does not deprotonate to yield an anionic state at physiological pH (36). Together these findings demonstrate the critical role of an anionic triazole in MESH1 inhibition.

Other tested analogs containing single or double halogenated substitutions at the benzene ring or alkylation at the triazole ring are shown in **(Supplemental Figure 1E)**. None of these compounds showed better activity than TBB, further supporting the importance of multiple halogens and an anionic triazole group. These results suggest that small structural changes to TBB derivatives significantly alter potency towards MESH1, underscoring the structural specificity of TBB in targeting MESH1.

The MESH1-TBB crystal structure and SAR data provide corroborative evidence of how TBB inhibits MESH1. Our structure provides a molecular explanation for the importance of the anionic state of TBB: The negative charge would result in an enhanced π-cation interaction with Arg-24. This stronger interaction helps explain why TBB inhibits MESH1, but the methylated analogs do not. The structure also explains why bromine may be the optimal halogen. The hydrophobic character of the bromine allows TBB to form hydrophobic interactions with the hydrophobic binding pocket. Iodine might be able to form stronger hydrophobic interaction. However, it appears that the larger orbital of iodine at the 6-position would sterically clash with Tyr-146. Collectively, the SAR—and its agreement with the co-crystal structure —defines a minimal pharmacophore (anionic 1,2,3-triazole plus 4,5,6,7-bromination) required for productive MESH1 engagement.

### TBB Protects Cells from Ferroptosis

After establishing the SAR of TBB, we investigated if MESH1 inhibition by TBB could protect cells from ferroptosis. During assay optimization, we noticed that TBB-mediated ferroptosis protection was attenuated under standard serum conditions. Since serum interactions have been reported for other benzotriazole compounds, we reasoned that TBB might bind to the 10% Fetal Bovine Serum (FBS) present in the cell culture media, leading to reduced activity (37). Consistently, incubation of TBB with serum resulted in substantial retention of TBB in a spin filter assay, and lowering serum concentrations enhanced ferroptosis protection **(Supplemental Figure 2A, B).** As a result, all subsequent cell culture experiments were performed under low-serum conditions (0.5% FBS).

Ferroptosis protection experiments were first performed using an RCC4 cell line, where ferroptosis was induced with erastin, and a serial dilution of TBB was added concurrently. Cell viability was assessed after 24 hours using the CellTiter Glo assay (Promega). TBB provided dose-dependent ferroptosis protection, consistent with *MESH1* genetic knockdown **(Figure 4A)** (14). TBB at 10 and 20 µM provided nearly complete protection against ferroptosis-induced cell death. TBB similarly protected HT-1080 and MDA-MB-231 cells from ferroptosis-induced cell death, suggesting that the mechanism of ferroptosis protection was generalizable across cell lines **(Supplemental Figure 2C, D)**. The TBB-mediated ferroptosis protection was further confirmed using CellTox Green assay (Promega), which monitors cell death via membrane rupture **(Figure 4B, C)**. To determine whether TBB regulates lipid peroxidation, RCC4 cells were incubated with or without TBB and erastin. Lipid peroxidation was monitored via C11 BODIPY Dye and flow cytometry. This experiment revealed that TBB significantly reduced lipid peroxidation in the cellular membrane **(Figure 4D, E)**. Because many ferroptosis-suppressing small molecules act as radical-trapping antioxidants, we additionally tested whether TBB exhibited intrinsic radical scavenging activity using the DPPH assay, a cell-free colorimetric assay that measures direct quenching of a stable free radical and observed ≤3% DPPH scavenging by TBB. This indicates no appreciable radical-trapping activity of TBB, in contrast to the robust activity of the positive control Trolox **(Supplemental Figure 2E)**. We next tested whether TBB acts through another common non-specific mechanism—iron chelation. We found that TBB did not prevent Fe²⁺–ferrozine complex formation in a ferrozine competition assay, in contrast to the strong chelation seen with deferoxamine **(Supplemental Figure 2F)**. Collectively, these experiments establish TBB as a specific ferroptosis suppressor whose activity is independent of radical-trapping or iron chelation.

**Figure 4.**
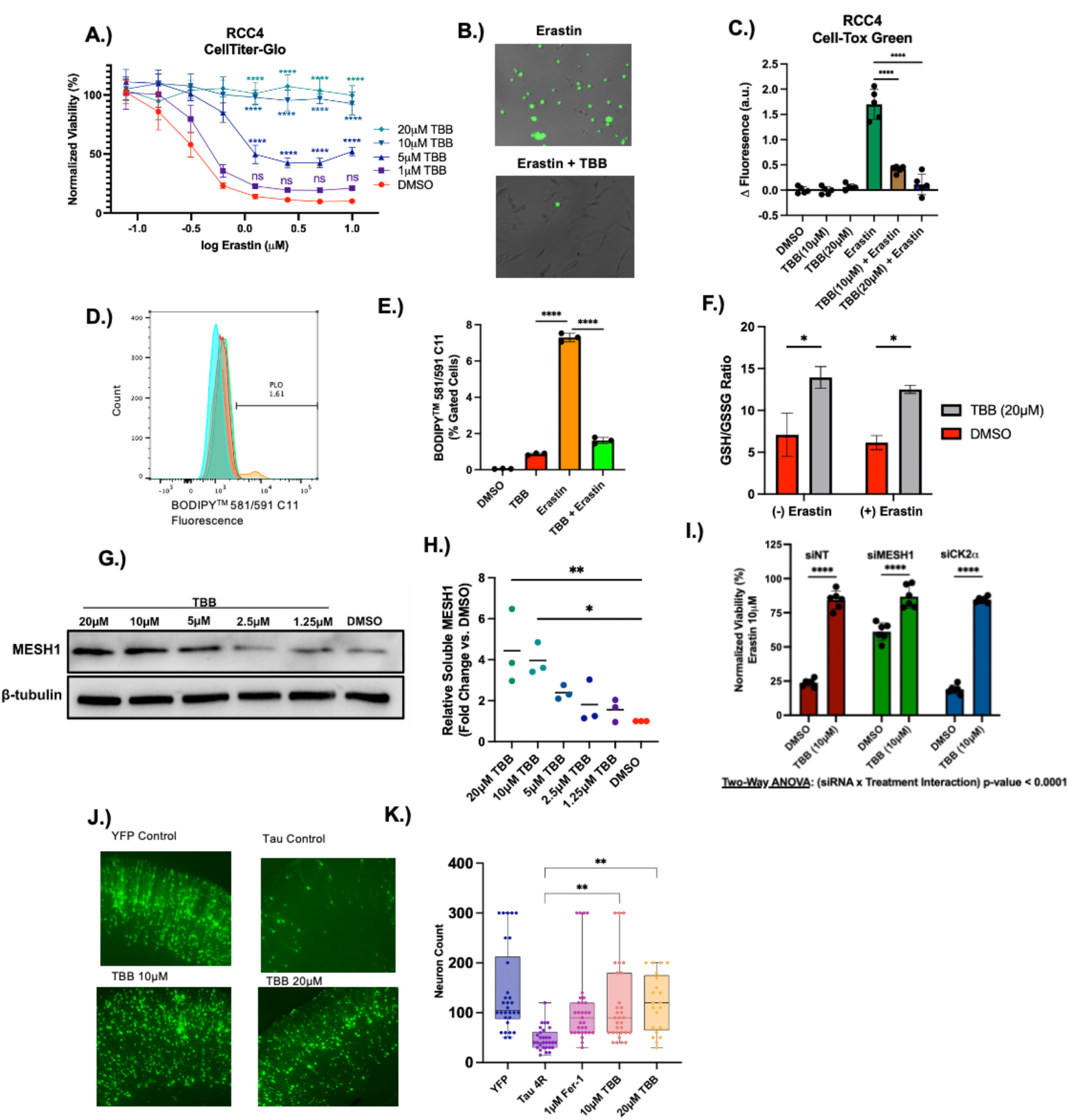
**A.** RCC4 cells were treated with the combination of erastin and TBB for 24hours. Cell viability was measured using CellTiter Glo assay. n=6, error bars = SD. Statistical significance shown at selected erastin doses was determined using Two-Way ANOVA with Dunnett’s multiple comparison of DMSO control vs. TBB concentration**. B.** Fluorescent microscopy of RCC4 cells incubated with erastin (0.625 µM) and TBB (20 µM) after 24 hours with inclusion of CellTox Green reagent. **C**. Quantification of CellTox Green fluorescent signal with plate reader of RCC4 cells treated with TBB and erastin (0.625 µM) for 24hours. n=5, error bars = SD. Statistical significance was determined with One-Way ANOVA with Dunnett’s multiple comparison of erastin vs. erastin + TBB samples. **D-E.** Measuring lipid peroxidation via C11-BODIPY^TM^ staining of RCC4 cells treated with or without TBB (20 µM) and erastin (5 µM) for 16 hours. Quantification of lipid peroxidation shown in **E**. n=3 error bars = SD. Statistical significance was determined with One-Way ANOVA with Dunnett’s multiple comparison of erastin vs. TBB and erastin vs erastin + TBB samples **F.** Measurement of GSH/GSSG ratio of RCC4 cells treated with or without erastin (1.25 µM) and with or without TBB (20 µM) for 6 hours. n=4 error bars = SEM. Statistical significance was determined with Two-way ANOVA with Sidák multiple comparisons (TBB vs. DMSO) **G.** Representative immunoblot showing soluble MESH1 and β-tubulin after treating RCC4 cells with a serial dilution of TBB concentrations (20 µM-1.25 µM) or DMSO control and heating the cells to 50 ℃ for 3 min. **H.** Quantification of relative fold increase of soluble MESH1. The MESH1 band intensity was normalized to β-tubulin loading control and then normalized to DMSO. n=3 independent experiments. Statistical Significance was assessed with One-Way Anova with Dunnett’s multiple comparisons test for comparing each TBB concentration to DMSO. **I.** Viability determined with CellTiter Glo of RCC4 cells transfected with siNT, siCK2⍺, or siMESH1 and treated with 10 µM erastin and DMSO or 10 µM TBB for 24 hours. n=6, error bars = SD. Statistical significance was determined using Two-Way ANOVA with Dunnett’s multiple comparison of DMSO vs. TBB. **J.** Representative images of neurons transfected with YFP+ or Tau and treated with TBB. **K**. Quantification of neuron survival. One-way ANOVA with Dunnett’s post-hoc test vs. Tau 4R. * = p-value <0.05, ** = p-value < 0.01, *** = p value <0.001, **** p-value < 0.0001

To better understand TBB’s role in suppressing ferroptosis, we further assessed whether TBB regulates the glutathione antioxidant defense system. The levels of free GSH and oxidized GSSG were quantified using the GSH/GSSG-Glo assay (Promega). Under conditions with or without erastin, TBB significantly increased the level of reduced glutathione while maintaining the level of oxidized glutathione, resulting in a higher GSH/GSSG ratio **(Figure 4F)**. Decreasing the cellular GSH/GSSG ratio impairs cellular antioxidant defenses, thereby increasing lipid peroxidation. Consistent with genetic knockdown of *MESH1*, TBB can both increase the GSH/GSSG ratio and reduce lipid peroxidation.

To determine if TBB directly engages with MESH1 in cells, we performed an isothermal Cellular Thermal Shift Assay (iCETSA). RCC4 cells were treated with a serial dilution of TBB and heated to 50 ℃, which resulted in the aggregation of MESH1. Compounds that bind to MESH1 can stabilize the protein and result in greater levels of soluble MESH1, which can be measured by immunoblot **(Figure 4G)**. TBB stabilized MESH1 in a dose-dependent manner, consistent with the biochemical IC_50_ and cellular ferroptosis protection phenotype data. At concentrations of 10 and 20 µM, TBB had a ∼4-fold increase in soluble MESH1 in comparison to the DMSO control **(Figure 4H)**. These findings provide strong evidence that TBB directly engages with MESH1 within cells.

After establishing cellular engagement of TBB with MESH1, we examined whether inhibition of MESH1, but not other TBB targets, underlies the ferroptosis-protection capabilities of TBB. We first sought to determine if CK2 inhibition could contribute to ferroptosis protection by individually knocking down *CK2α* and *MESH1* with siRNA in HT-1080 cells **(Supplemental Figure 2G).** We found that only knockdown of *MESH1* resulted in ferroptosis protection. To further evaluate the contribution of MESH1, RCC4 cells were transfected with non-targeting, *CK2α* or *MESH1* targeting siRNA and treated with erastin in the presence or absence of TBB **(Figure 4I, Supplemental Figure 2H)**. TBB significantly increased cell viability across all conditions. Notably, the dynamic range of TBB rescue was considerably reduced in *MESH1* depleted cells compared to siNT or *CK2α* knockdown. This attenuation is consistent with the notion that TBB mediated MESH1 inhibition protects against ferroptosis. The persistence of TBB rescue in *MESH1* knockdown likely reflects incomplete knockdown (∼80% reduction, **Supplemental Figure 2I**) and/or engagement with additional pathways that modulate ferroptosis sensitivity. Together, these results indicate that CK2 does not play a substantial role in ferroptosis protection and highlights an important role of TBB inhibition of MESH1 in ferroptosis protection.

To evaluate the ferroptosis-protective effect of TBB in a disease-relevant context, we used an ex vivo rat hippocampal slice system for Alzheimer’s disease (AD), in which Tau and YFP were introduced by gene gun transfection. Canonical ferroptosis inhibitors were found to inhibit neuronal death (38, 39), suggesting that ferroptosis is an important contributor to these AD models. Neuron viability was assessed with fluorescence over the course of three days **(Figure 4J, K)**. The brain slices were treated with vehicle control and TBB. Ferrostatin-1, a ferroptosis-suppressing RTA, was included as a positive control. Treatment with TBB increased neuronal survival compared to vehicle control. The positive control, ferrostatin-1, also protected neurons, highlighting the potential role of ferroptosis in Tau pathology. Collectively, our data show that TBB protects against ferroptosis in cells and in an ex vivo model of Alzheimer’s disease.

## Discussion

In this study, we have established TBB as a small-molecule inhibitor of MESH1, defined its key binding determinants through co-crystal structure analysis and SAR, and demonstrated on-target suppression of ferroptosis across multiple cell lines and in an ex vivo neuronal model of AD.

Unlike nonspecific radical-trapping antioxidants that are limited by stoichiometry, inhibition of MESH1 preserves endogenous NADPH-dependent antioxidant pathways, offering a mechanistically distinct route to ferroptosis control.

The co-crystal structure of the MESH1-TBB complex reveals that TBB occupies the nucleotide-binding site of MESH1 in a manner resembling the binding of the adenine moiety of NADPH. Our structure reveals key interactions between TBB and MESH1 that rationalize the observed SAR of TBB analogs and explain why certain TBB derivatives exhibit markedly reduced potency. Although TBB has traditionally been considered a selective CK2 inhibitor, our data indicate that its ferroptosis-related cellular phenotypes—including protection from ferroptosis-induced cell death, decreased lipid peroxidation, and elevated GSH/GSSG ratios—are mediated by MESH1 inhibition. Consistent with this conclusion, *CK2α* knockdown does not confer protection against ferroptosis. Moreover, prior studies have shown that CK2 activity supports SCD1-dependent oleic acid production, which mitigates ferroptosis (40–42). Accordingly, direct inhibition of CK2 by TBB would be expected to promote ferroptosis, a prediction that is inconsistent with the observed protective effect of TBB. Finally, iCETSA confirms engagement of MESH1 by TBB in cells, and the protective phenotype is abrogated upon *MESH1* knockdown, establishing MESH1 as the functional target underlying TBB-mediated ferroptosis inhibition.

Our study demonstrates the feasibility of developing MESH1-targeting small-molecule inhibitors for therapeutic suppression of ferroptosis in cell lines and ex vivo neurodegenerative disease models. Although the benzotriazole core of TBB exhibits activity against multiple kinases in profiling assays—including CK2, PIM, and HIPK family members (43)—our genetic and biophysical data indicate that cellular engagement of MESH1 is required for the TBB-mediated protection from ferroptosis. Future medicinal chemistry efforts will focus on improving the specificity and potency of TBB derivatives toward MESH1. Notably, our co-crystal structure reveals that TBB occupies only a small portion of the MESH1 active site, highlighting substantial opportunities for future structure-guided optimization.

The small size and hydrophobic character of the TBB scaffold are consistent with compounds capable of penetrating the blood-brain barrier. Consistently, TBB has been administered in rodent models and shown to modulate central nervous system phenotypes, indicating that the scaffold can access the brain (44, 45). Our *ex-vivo* brain slice model demonstrated that TBB and other ferroptosis inhibitors could protect neurons from cell death in an AD model. Importantly, CK2 activity has also been implicated in Alzheimer’s disease related pathways including signaling upstream of Tau phosphorylation and inhibition of CK2 has been shown to promote neuron survival (46–49). Additionally, CK2 has been reported to be upregulated in patients with AD (47). A future MESH1 inhibitor with increased specificity would allow for the dissection of MESH1 ferroptosis protection properties and CK2 kinase activity in the context of tau pathology. Advances would enable more comprehensive *in vivo* studies and determine whether targeting MESH1 is a viable strategy for neurodegenerative therapeutics.

In conclusion, this study establishes TBB as the first reported small molecule inhibitor of MESH1 and defines a structural and biochemical framework for targeting MESH1 to suppress ferroptosis. Unlike traditional RTAs, MESH1 inhibition by TBB offers a distinct strategy for ferroptosis protection by preserving the endogenous NADPH pool. Our findings position MESH1 as a tractable and mechanistically compelling regulator of ferroptosis for therapeutic intervention. Given the involvement of ferroptosis in diseases such as neurodegeneration, future development of selective MESH1 inhibitors may enable therapeutic approaches for ferroptosis-driven pathologies.

## Methods

### Chemicals

Erastin (MedChem Express:HY-15763), IKE (MedChem Express: HY-114481), TBB (MedChem Express: HY-14394), TFBt (Enamine; EN300-6487228), TCBt (Sigma; COMH04236427), TIBi (AABlocks; AA01X6YQ), 5-monoBBt (Enamine; EN300-137666), 4-monoBBt (Enamine; EN300-137659), 5,6-diBBt (Enamine; EN300-6775516), 5-monoIBt (Enamine; EN300-2951206), 4-F,5-monoBBt (Enamine; EN300-5495791), 6,7diBQX (Enamine; EN300-19483685), DMAT (MedChem Express: HY-15535), TBCA (MedChem Express: HY-110052), DRB (MedChem Express: HY-14392), TBBi (Cayman Chemical, 18886), DPPH (Thermo Scientific, 044150.MD), Trolox (Thermo Scientific, 218940010), Ferrozine (Sigma, 160601), DFO (Sigma, D9533)

### Synthesis Procedures and Analytical Data

#### Preparation of N^1^-EtNH_2_-TBBt

*N^1^*-EtNH_2_-TBBt was synthesized according to a previously reported procedure(50). Spectral data of *N^1^*-EtNH_2_-TBBt were identical to those reported(50). *N^1^*-EtNH_2_-TBBt: ^1^H NMR (700 MHz, CDCl_3_) δ 5.01 (t, J = 6.3 Hz, 2H), 3.33 (t, J = 6.2 Hz, 2H); HRMS (ESI-TOF) m/z: [M + H]^+^ Calcd for C_8_H_7_Br_4_N_4_ 474.7398; Found 474.7395.

#### Preparation of N^1^-Me-TBBt and N^2^-Me-TBBt

*N*^1^-Me-TBBt and *N*^2^-Me-TBBt were synthesized according to a previously reported procedure(51). Spectral data of *N*^1^-Me-TBBt and *N*^2^-Me-TBBt were identical to those reported(51). For *N*^1^-Me-TBBt: ^1^H NMR (700 MHz, CDCl_3_) δ 4.56 (s, 3H); ^13^C NMR (176 MHz, CDCl_3_) δ 145.55, 132.26, 129.03, 124.41, 116.70, 105.92, 38.17. For *N*^2^-Me-TBBt: ^1^H NMR (700 MHz, CDCl_3_) δ 4.55 (s, 3H); ^13^C NMR (176 MHz, CDCl_3_) δ 143.38, 126.48, 113.64, 44.08; HRMS (ESI-TOF) m/z: [M + H]^+^ Calcd for C_7_H_4_Br_4_N_3_ 445.7133; Found 445.7126

#### Preparation of 4,5,6-tribromobenzotriazole

4,5,6-Tribromobenzotriazole was synthesized according to a previously reported procedure(35). Spectral data of 4,5,6-tribromobenzotriazole were identical to those reported(35).^1^H NMR (700 MHz, DMSO) δ 8.46 (s, 1H); HRMS (ESI-TOF) m/z: [M + H]^+^ Calcd for C_6_H_3_Br_3_N_3_ 353.7872; Found 353.7867

### MESH1 Purification

Recombinant human MESH1 was purified as previously described (14). Briefly, MESH1 was cloned into a modified pET28A plasmid containing a His-10-Sumo tag at the N-Terminus of MESH1. The plasmid was transformed into BL21 cells and grown at 37°C, 200 RPM to an OD600 nm of 0.5 and expression was induced with the addition of 1mM isopropyl β-d-1-thiogalactopyranoside (GoldBio) for 2 hours. Cells were lysed using the French Press at 1200 PSI and the protein was purified using Ni Affinity chromatography (Cytivia). The His-10-Sumo tag was cleaved using a SENP1 protease and MESH1 was further purified using a second round of Ni Affinity Chromatography. Recombinant MESH1 was further purified using size exclusion chromatography with Superdex 75 resin with a buffer containing 200mM NaCl, 50mM Tris pH8, and 0.1% beta-mercaptoethanol.

### Library Screening and IC_50_ Calculation

For Screening of the Cayman Comprehensive Kinase Library, MESH1 was mixed with NADPH and Inhibitor from library at final concentrations of 100 nM MESH1, 200 µM NADPH, 10 µM inhibitor in a buffer containing 200 mM NaCl, 50 mM Tris pH8, and 1 mM MnCl_2_. The reaction was run at 37°C for 10 minutes and was quenched with formic acid. The level of phosphate released was quantified using Malachite Green Assay (Sigma, MAK-307), according to the manufacturer’s instructions with 3:1:1 ratio of Formic Acid: Enzyme/Inhibitor reaction: Malachite Green Assay Working solution. Phosphate was quantified by measuring absorbance at 620nm.

For calculating percent activity of TBB analogs, a similar procedure was performed using 75 nM MESH1 110 µM NADPH, 100 µM inhibitor in a buffer containing 200 mM NaCl, 50 mM Tris pH 8, and 1 mM MnCl_2._ The reaction was run at 37°C and quenched after 10 minutes. Data was collected with n=3 replicates, +/- standard deviation.

A similar procedure was used to calculate the IC_50_ value of inhibitors. IC_50_ values were calculated using 18 data points of two-fold serial dilutions of inhibitor ranging from 500 to 0.0038 µM. MESH1 protein was mixed with NADPH and Inhibitor at final concentrations of 100 nM MESH1, 110 µM NADPH and serial dilution of inhibitor. Data points were collected at 1,2,4,5, and 6 min and the reaction was quenched in 5 M Formic Acid. Phosphate release was calculated using Malachite Green Assay (Sigma, MAK-307). Vi/Vo was calculated by dividing the velocity of inhibitor sample with the velocity of n=4 positive control samples lacking inhibitor. The IC_50_ values were calculated by plotting Vi/Vo values against the log (inhibitor concentration) and fitting with four parameter non-linear regression fitting using GraphPad Prism software. Data was collected with n=3 independent experiments, +/- standard deviation. Apparent inhibition constants K_i_^app^ were calculated from IC50 values using the Cheng-Prusoff equation assuming competitive inhibition.

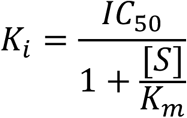

### Thermal Shift Assay

Thermal shift assays were performed using a StepOnePlus Real Time PCR System (Applied Biosystems). Reactions of 20 µl containing 3 µM MESH1, 5X Sypro Orange Dye (Sigma, S5692) 3% DMSO with/without 100 µM Inhibitor and a buffer consisting of 50 mM Tris pH 8, and 200 mM NaCl. Samples were loaded into qPCR Plate and heated from 25°C to 95°C using a 2% ramp rate and fluorescence was continuously monitored through the SYBR channel. Melting Temperatures-Tm were calculated from the first derivative of fluorescence curve using StepOne Software (v2.2.2) and ΔTm was calculated relative to DMSO control.

### MESH1 Inhibitor Crystallography

For the MESH1-TBB structure, Apo-MESH1 crystals were generated using sitting drop vapor diffusion method. MESH1 was concentrated to 9 mg/ml and mixed 1:1 with a mother liquor solution containing 14% PEG 3350, 0.2 M Malic Acid pH 7, and 0.1 M Tris pH 7.5 and the crystals were harvested after growing for 10 days at 20°C. TBB was dissolved into PEG 300 and soaked into the Apo-MESH1 crystals. A soaking/cryo-protectant solution was prepared containing 3:2 mother liquor: cryo-protectant containing final concentrations of 20% glycerol, 16% PEG 300, 6 mM TBB. Crystals were soaked for 2 hours.

X-Ray Diffraction data was collected at the Northeastern Collaborative Access Team (NECAT) 24-ID-E beamline at the Advanced Photon Source at Argonne National Laboratory. The diffraction data was processed using the XDS package (52). The phase data of the MESH1 Inhibitor complex was prepared through molecular replacement using the PHASER program in PHENIX, using PDB ID: 5VXA as the input (53). Restraints for the inhibitor were generated using Grade2. Iterative model building was performed using COOT (54) and Phenix(53). Figures were generated using Pymol. **Δ**SASA was calculated using PDBePISA EMBL-EBI webserver(55). Protein ligand interactions were calculated and visualized with PLIP(56).

### TBB-FBS Binding Experiment

TBB (100 µM, 5% DMSO) was incubated at room temperature with phosphate buffered saline (PBS) containing 10%,1%,0.1%, 0% FBS for an hour. Unbound TBB was quantified by passing the solution through a 3,000 MWCO (Amicon)spin filter and measuring absorbance λ= 290 nm using a NanoDrop One (ThermoFisher).

### Cell Culture

RCC4, HT-1080, and MDA-MB-231 cells were obtained from Duke University’s Cell Culture Facility. The cell lines were authenticated with STR DNA profiling and verified to be free from mycoplasma contamination. The cell lines were maintained in a humidified incubator at 37°C with 5% CO_2_. Prior to experiments, the cells were cultured in DMEM (GIBCO-11995-065) with 10% heat-inactivated FBS (ThermoFisher: 10082147), and 1% Streptomycin (10,000 µg/ml)/Penicillin (10,000 Units/ml) (GIBCO-15140-122).

### Cell Viability and GSH/GSSG Experiments

Cell viability was monitored using CellTiter-Glo (Promega) assay according to the manufacturer’s instructions. Briefly, cells were plated 3,000 cells/well (RCC4) or 5,000 cells/well (HT-1080/MDA-MB-231) in a media consisting of DMEM (GIBCO-11965-092), 0.5% FBS, 1% Pen/Strep. After 24 hours, erastin/IKE and TBB were added at concentrations at which the DMSO percentage was held constant and did not exceed 0.5%. After 24 hours, the 96-well plates were incubated at room temperature for half an hour and 20 µl of CellTiter-Glo working reagent was added to 100 µl of media. After 15 minutes of continuous shaking, luminescence was recorded with a Varioskan Lux plate reader (ThermoFisher Scientific).

For quantification of cell death, CellTox Green (Promega) assay was used according to manufacturer’s instructions. RCC4 cells were plated 3,000 cells/well in a black 96-well plate with media consisting of DMEM (GIBCO-11965-092), 0.5% FBS, 1% Pen/Strep. After 24 hours, TBB (0,10, or 20 µM) and erastin (0 or 0.625 µM) were added to the plate with a DMSO percentage of 0.5% and CellTox Green Reagent dilution to a working concentration of 1X. After 24 hours, cells were assessed with microscopy and a Varioskan Lux plate reader (ThermoFisher Scientific) recording fluorescence (excitation: 485nm and emission: 520nm).

To quantify GSH/GSSG levels, the GSH/GSSG-Glo (Promega) assay was used. RCC4 cells were plated 3,000 cells/well in a 96 well plate in consisting of DMEM (GIBCO-11965-092), 0.5% FBS, 1% Pen/Strep. After 24 hours, TBB (0 or 20 µM) and erastin (0 or 1.25 µM) with a DMSO percentage of 0.5%. After six hours of incubation with the drugs, GSH and GSSG were calculated with the kit according to the manufacturer’s instructions. Briefly, the cells were lysed using total or oxidized GSSG lysis reagent. Afterwards, luciferin generation reagent was added followed with the addition of luciferin detection reagent. Luminescence was recorded with a Varioskan Lux plate reader (ThermoFisher Scientific) and GSH/GSSG levels were quantified from luminescence measurements.

### Flow Cytometry

For lipid peroxidation staining, 1.5 x 10^5^ RCC4 cells were added to a 6-well plate for 24 hours. Next, TBB (0 or 20 µM) and erastin (0 or 5 µM) were added to the cells with a DMSO percentage of 0.5%. After 16 hours, the cells were washed with PBS and incubated with 10 µM of BODIPY™ 581/591 C11 (Invitrogen, D3861) diluted in DMEM media and the cells were incubated for 1 hour at 37°C. After a subsequent PBS wash, the cells were resuspended with 0.25% trypsin EDTA washed with PBS and filtered into a flow cytometry tube. Lipid peroxidation was quantified through flow cytometry analysis BD FACSCanto^TM^ II (Biosciences).

### DPPH Radical Scavenger Assay

DPPH (2,2-Diphenyl-1-picrylhydrazyl) was prepared in methanol and mixed 1:1 with a serial dilution of Trolox/TBB, also prepared in methanol, in a clear 96-well plate for final concentrations of 100 µM DPPH, 100-3.125 µM Trolox/TBB, and DMSO at 0.5%. The plate was incubated in the dark for 30 min and Absorbance at 517 nm was recorded using a Varioskan Lux plate reader (ThermoFisher Scientific). Each measurement was blank subtracted and % DPPH radical scavenging activity was calculated by comparing sample to vehicle control.

### Ferrozine Competition Assay

Compounds (TBB, DFO) or Vehicles control were prepared in a sodium acetate buffer (pH 5.5) and mixed with freshly prepared Fe^2+^ for final concentrations of 100 µM TBB/DFO and 50 µM Fe^2+^ with DMSO at 2%. The samples were incubated for 10 minutes prior to the addition of ferrozine at a final concentration 100 µM. After an additional 10 minutes of incubation Absorbance at 562 nm was measured using a Varioskan Lux plate reader (ThermoFisher Scientific) to quantify the ferrozine-Fe^2+^ complex. Each measurement was blank subtracted with its corresponding compound alone sample.

### Isothermal Cellular Thermal Shift Assay (CETSA)

Isothermal CETSA was performed to assess the target engagement of TBB with MESH1 in intact cells, according to a previously established protocol (57). RCC4 cells were grown to 80% confluency in one 10cm-dish (one per condition). The cells were treated with TBB at concentrations ranging from 20 to 1.25 µM (2-fold serial dilution of TBB) or 0.5% DMSO control at 37°C for an hour. The cells were resuspended in PBS containing inhibitor and cOmplete protease inhibitor (Roche) and harvested by gently scraping. Cells suspensions were incubated at room temperature for 3 minutes before heating at 50°C for 3 minutes. To lyse the cells, the samples underwent three snap-freeze/thaw cycles using liquid nitrogen. The samples were spun down 16,000xg for 35 minutes at 4°C and the supernatant was further clarified with an additional 16,000xg for 35 minutes at 4°C. Protein concentration was measured using a BCA assay (ThermoFisher) and equal amounts of protein were mixed with 4X SDS loading buffer and run on a 4-20% gradient SDS PAGE gel followed by immunoblotting. Proteins were transferred to 0.2 µm PVDF membranes and the membranes were cut horizontally at ∼37kDa with the upper portion used to probe β-Tubulin and the bottom portion to probe quantify MESH1. Membranes were blocked in 5% BSA and then incubated at 1:1000 overnight at 4°C with anti-β-Tubulin (Cell-Signaling Technology) or anti-MESH1 (Protein-Tech), followed by HRP-conjugated secondary antibodies treated 1:3000 at room temperature for an hour. The blots were developed using Amersham ECL reagent (GE) and imaged using ChemiDoc System (BioRad). The intensity of bands were quantified using FIJI (ImageJ), normalized to DMSO control and plotted as relative-fold change in MESH1 levels.

### siRNA Gene Silencing

Gene knockdown was performed using the following siRNA: siNT-AllStars Negative Control siRNA (Qiagen, SI03650318), siMESH1 (Target sequence: GGGAAUCACUGACAUUGUG, D-031786-01, Dharmacon), siCK2α (Target sequences: GAUCCACGUUUCAAUGAUA, GAUGUACGAUUAUAGUUUG, ACUGCUUGCUGGUCGCUUA, GGGCAGACACUCUCGAAAG, M-003475-03, Dharmacon). Knockdown was validated by quantitative RT-PCR. For viability assays, 2,500 RCC4 cells per well or 3,000 HT-1080 cells per well were reverse transfected with 2.5 pmol of siRNA and 0.15 µl of RNAimax (Thermo Scientific, 13778150) per well with a total volume of 100 µl. After 48 hours, the inhibitors were added at described concentrations and cellular viability was measured 24 hours later.

### Quantitative Real-Time PCR

RNA was extracted using the RNeasy Mini Kit (Qiagen, #74104) with DNase I treatment (Qiagen, #79524). cDNA was synthesized from 700 ng of total RNA with or without reverse transcriptase, which was prepared using SuperScript II reverse transcriptase (Thermo Scientific, 18064014) with random hexamers. Quantitative real-time PCR was performed using a StepOnePlus Real Time PCR System (Applied Biosystems) instrument and using Power SYBR Green PCR Master Mix (Thermo Scientific, 4368577). Primers used in the qRT-PCR were as follows: CK2α (Forward: GGTGAGGATAGCCAAGGTTCTG; Reverse: TCACTGTGGACAAAGCGTTCCC), MESH1 (Forward: GAGGCGGGAATCACTGACATTG; Reverse: GAGTCTTGTCATCTGTTACCTCC) and GAPDH (Forward: GTCTCCTCTGACTTCAACAGCG; Reverse: ACCACCCTGTTGCTGTAGCCAA).

### Brain Slice AD Model

Preparation of hemi-coronal brain slice explants from postnatal day 8 (P8) CD Sprague Dawley rats were prepared and maintained in neurobasal medium (which used NB 15 mM KCl + MK 801; Neurobasal A medium supplemented with 15% heat-inactivated pig and rat serum, 10 mM KCl, 10 mM HEPES, 100 U/ml penicillin/streptomycin, 1 mM sodium pyruvate, and 1 mM L-glutamine set in 0.5% reagent-grade agarose). Under sterile conditions, brains were dissected and cut into 250 microns hemi-coronal slices on a vibratome in chilled culture medium. Brain slices were plated into 12-well plates in interface configuration atop a solid culture medium made by adding 0.5% agarose with DMSO control or Compound. After explanting the brain slices, plates were placed for recovery at 30°C for 30 minutes in a humidified incubator under 5% CO_2_. After which, biolistic transfection of YFP and tau mutant 4R0N (tau4R) was done by transfecting YFP along with an empty vector or transfecting YFP along with tau 4R0N (all constructs in gWiz backbone). Slice cultures were maintained at 30 °C for 72 hours in a humidified incubator under 5% CO_2_. Brain Slice visualization was done using Leica MZIIIFL fluorescence stereomicroscope. Neuronal/Cortical counts were performed manually on day 3.

## Acknowledgements

This work was supported by NIH grants R01GM124062 (to J.T.C. & P.Z.) and R21AG077075 (to J.T.C. & J.H.). This work is based upon research conducted at the Northeastern Collaborative Access Team beamlines, which are funded by the National Institute of General Medical Sciences from the National Institutes of Health (P30 GM124165). The Eiger 16M detector on the 24-ID-E beamline is funded by a NIH-ORIP HEI grant (S10OD021527). This research used beamtime awards (DOI: https://doi.org/10.46936/APS-191720/60015447) from the Advanced Photon Source, a U.S. Department of Energy. Office of Science User Facility operated for the DOE Office of Science by Argonne National Laboratory under Contract No. DE-AC02-06CH11357.

## Author Contribution

The experimental strategy was conceived by J.H., P.Z., and J.-T.C. A.A.M. designed and performed biochemical, structural, and cellular experiments. X-ray crystallography experiments were performed and analyzed by A.A.M. under the guidance of P.Z. Biochemical and cell-based ferroptosis assays were performed and analyzed by A.A.M., with contributions from J.W., under the guidance of J.H., P.Z., and J.-T.C. Y.O. synthesized compounds under the guidance of J.H. D.D. performed brain-slice experiments under the guidance of S.R.F. Y.S., S.Y.C., C.-C.L., P.J., G.N., C.S.C., C.M., and D.R. contributed to experimental design and data interpretation. A.A.M, J.H., P.Z., and J.-T.C. wrote the manuscript

## Data availability

The data that support the findings of this study are available from the corresponding authors upon request. The coordinates of the MESH1-TBB complex have been deposited in the Protein Data Bank under the accession code 9ZZ9.

