## Supplemental Figures for "Identification of 4,5,6,7-Tetrabromo-1*H*-benzotriazole (TBB) as a Small Molecule MESH1 Inhibitor that Suppresses Ferroptosis"

Supplemental Material

**Table S1. X-ray data collection and refinement statistics of the MESH1-TBB Complex**

| | **MESH1/ TBB (9ZZ9)** |
|---|---|
| **Data collection** | |
| Space group | C 2 2 21 |
| Cell dimensions | |
| *a*, *b*, *c* (Å) | 72.4 78.37 62.7 |
| α, β, γ (°) | 90.00, 90.00, 90.00 |
| Resolution (Å) | 40.56 - 2.33 (2.413 - 2.33) |
| $R_{merge}$ | 0.2105 (1.071) |
| $R_{pim}$ | 0.06048 (0.2961) |
| $CC_{1/2}$ | 0.998 (0.891) |
| $I / \sigma I$ | 13.77 (3.10) |
| Completeness (%) | 99.92 (100.00) |
| Redundancy | 13.2 (13.9) |
| **Refinement** | |
| Resolution (Å) | 2.33 |
| No. reflections | 7906 (783) |
| $R_{work}$ / $R_{free}$ | 0.2081/ 0.2277 |
| No. atoms | 1455 |
| Protein | 1410 |
| Ligand/ion | 14 |
| Water | 31 |
| *B*-factors | 46.40 |
| Protein | 46.38 |
| Ligand/ion | 56.56 |
| Water | 42.83 |
| R.m.s. deviations | |
| Bond lengths (Å) | 0.002 |
| Bond angles (°) | 0.39 |
| Ramachandran | |
| favored (%) | 99.43 |
| allowed (%) | 0.57 |
| outliers (%) | 0.0 |

*Values in parentheses are for highest-resolution shell.

A.) B.) C.)

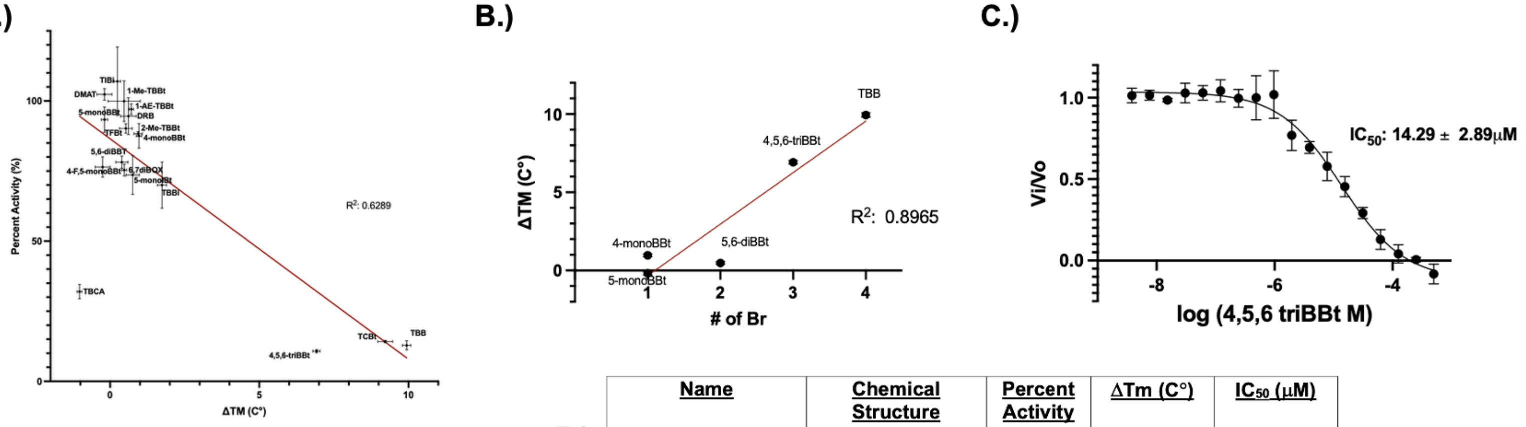


D.)

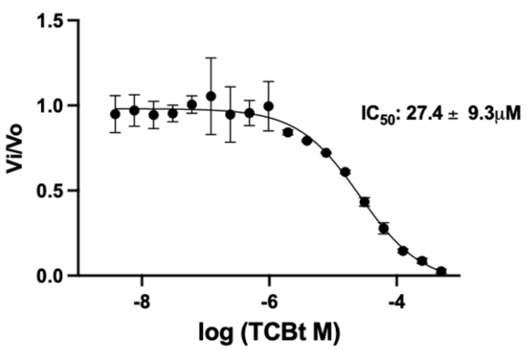


E.)

| Name | Chemical Structure | Percent Activity at 100μM (%) | ΔTm (C°) | $IC_{50}$ (μM) |
|---|---|---|---|---|
| TBB<br>4,5,6,7-tetrabromobenzotriazole | 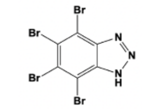 | 12.85 +/- 1.64 | 9.94 +/- 0.14 | 4.7 +/- 0.3 |
| 5-monoIBt<br>5-iodo-1H-1,2,3-benzotriazole | 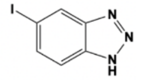 | 73.6 +/- 6.93 | 0.76 +/- 0.22 | - |
| 4-F,5-monoBBt<br>4-fluoro-5-bromo -1H-1,2,3-benzotriazole | 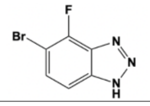 | 78.16 +/- 2.14 | 0.39 +/- 0.20 | - |
| 1-AE-TBBt<br>4,5,6,7-tetrabromo-1-(2-aminoethyl)benzotriazole | 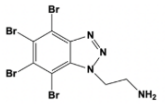 | 99.92 +/- 7.23 | 0.47 +/- 0.54 | - |
| 6,7diBQX<br>6,7-dibromoquinoxaline | 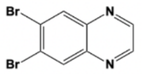 | 76.46 +/- 3.62 | -0.25 +/- 0.25 | - |

**Supplemental Figure 1. A.** Correlation of enzyme inhibition (percent activity) and thermal shift (**Δ**Tm) at 100µM inhibitor. Data fitted with linear regression (n= 3 for both experiments, mean ± S.D). **B.** Correlation of # or Br and thermal shift (**Δ**Tm). Data fitted with linear regression (n= 3, mean ± S.D). **C.** IC50 curve of 4,5,6-triBBt fitted with four-parameter fitting (n= 3 independent experiments, mean ± S.D.) **D.** IC50 curve of TCBt fitted with four-parameter fitting (n= 3 independent experiments, mean ± S.D.). **E.** Table of other TBB analogs tested. In heat map red color = stronger inhibition and larger thermal shift and white = weaker inhibition and smaller thermal shift.

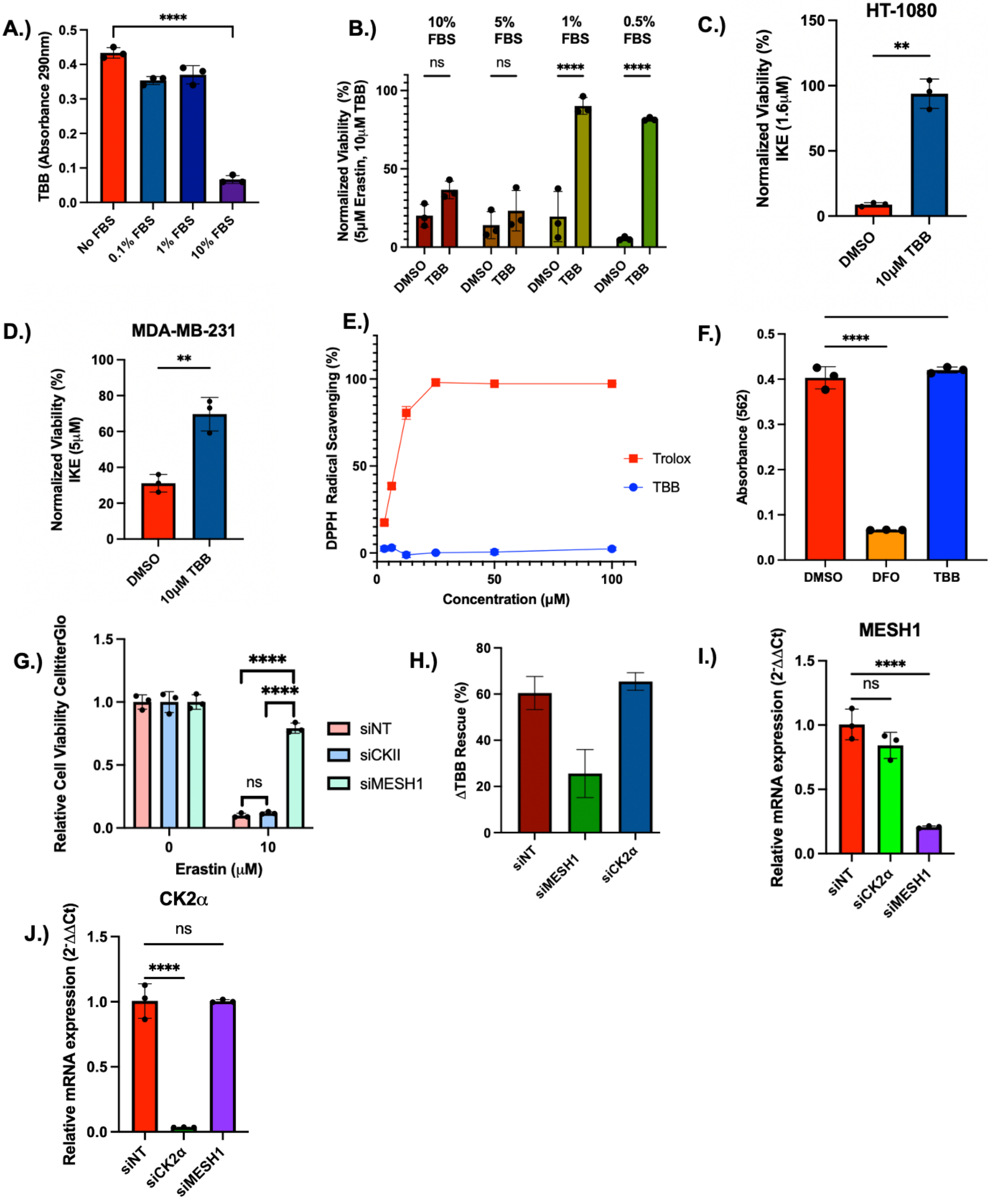


**Supplemental Figure 2. A.** Quantification of unbound TBB after incubation with various FBS concentrations. n=3 technical replicates, error bars = SD. One-way ANOVA with Dunnett's post-hoc test vs. DMSO **B.** RCC4 cells were treated 5 μM erastin and DMSO or 10μM TBB at the indicated fetal bovine serum (FBS) concentrations and cell viability was determined with CellTiterGlo and normalized to control. n=3, error bars = SD. Two-way ANOVA with Sidák multiple comparisons (DMSO vs. TBB) **C-D.** CellTiter Glo assessment of cellular viability of HT-1080 and MDA-MB-231 treated with the combination of IKE and 10 μM TBB. n=3, error bars = SD. Unpaired two-tailed T-test (DMSO vs. TBB) **E.** DPPH radical scavenging activity of TBB and Trolox across the indicated concentrations. n=4 technical replicates error bars =SD. **F.** DMSO, DFO(100 μM) and TBB (100 μM) were incubated with $Fe^{2+}$, prior to the addition of ferrozine. A562nm was measured to quantify free $Fe^{2+}$.n=3, error bars = SD. One-way ANOVA with Dunnett's post-hoc test vs. DMSO **G.** CellTiter Glo measuring cell viability of HT-1080 cells treated with siNT, siCK2, and siMESH1 with or without the treatment of 10μM erastin.**H**. Quantification of TBB mediated rescue. **I, J.** RCC4 cells were transfected with siNT, siCK2α, siMESH1 for 48hours and mRNA levels were quantified by RT-PCR using GAPDH for normalization. Relative expression was calculated by $2^{-}$ ΔΔCt method. n= 3 biological replicates, error bars= SD. One-way ANOVA with Dunnett's post-hoc test vs. siNT

* = p-value <0.05, ** = p-value < 0.01, *** = p value <0.001, **** p-value < 0.0001

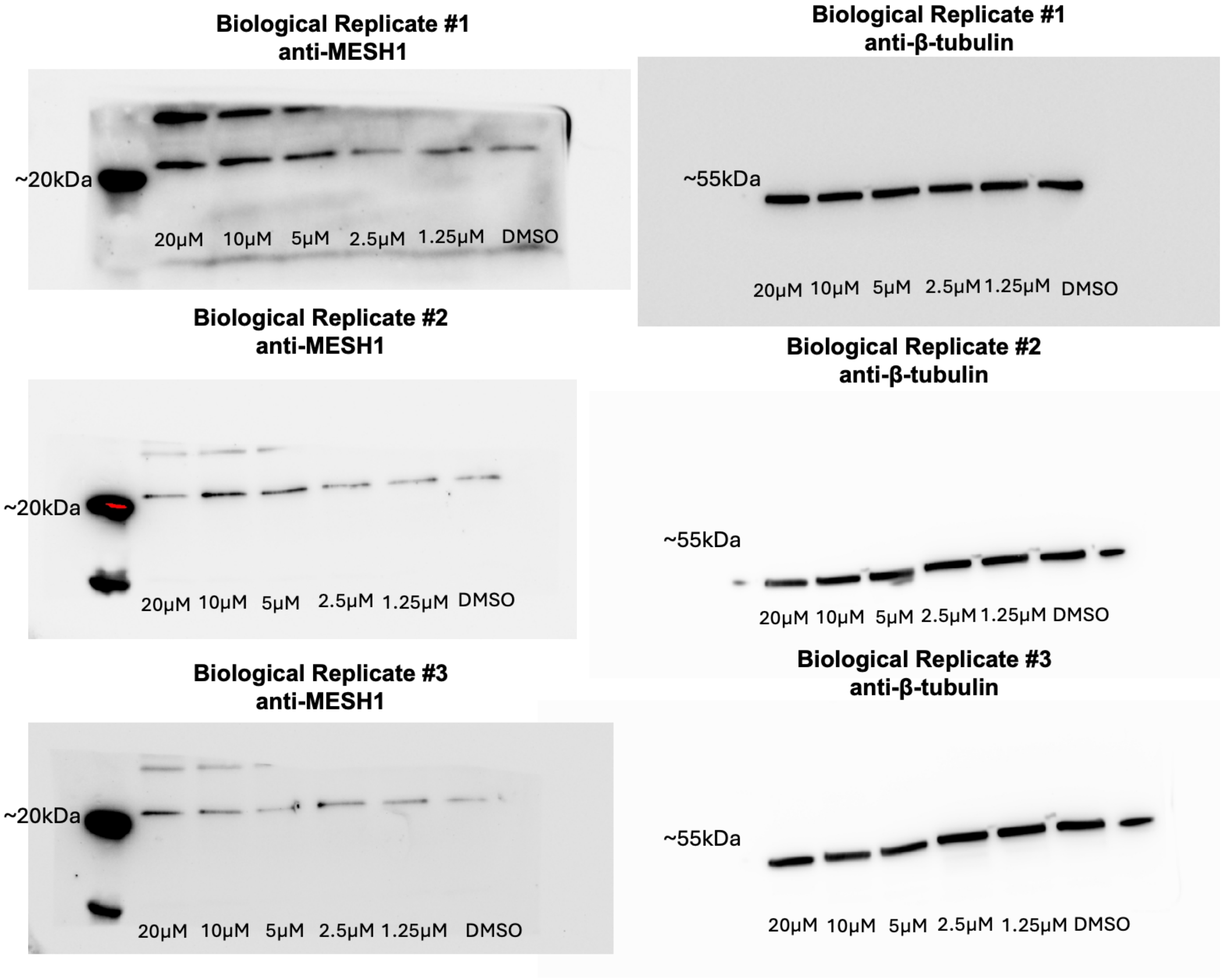


**Supplemental Figure 3.** Unprocessed Western Blots for Figure 4H. The MESH1 blot was captured at a longer exposure than the β-tubulin blot and the membrane outline of β-tubulin blots were not visible due to the short exposure.
